# DEFORMATION BASED MORPHOMETRY STUDY OF LONGITUDINAL MRI CHANGES IN BEHAVIORAL VARIANT FRONTOTEMPORAL DEMENTIA

**DOI:** 10.1101/670646

**Authors:** Ana L. Manera, Mahsa Dadar, D. Louis Collins, Simon Ducharme, Frontotemporal Lobar Degeneration Neuroimaging Initiative

**Author notes:** These authors contributed equally to this work (D. Louis Collins, Simon Ducharme). Data used in preparation of this article were obtained from the Frontotemporal Lobar Degeneration Neuroimaging Initiative (FTLDNI) database (http://4rtni-ftldni.ini.usc.edu/). The investigators at NIFD/FTLDNI contributed to the design and implementation of FTLDNI and/or provided data but did not participate in analysis or writing of this report (unless otherwise listed). **Address for correspondence:** D. Louis Collins, PhD, Montreal Neurological Institute, McConnell Brain Imaging Centre, 3801, University, Montreal, Quebec, H3A 2B4.

## Abstract

**Objective:** To objectively quantify how cerebral volume loss could assist with clinical diagnosis and clinical trial design in the behavioural variant of frontotemporal dementia (bvFTD).

**Methods:** We applied deformation-based morphometric analyses with robust registration to precisely quantify the magnitude and pattern of atrophy in patients with bvFTD as compared to cognitively normal controls (CNCs), to assess the progression of atrophy over one year follow up and to generate clinical trial sample size estimates to detect differences for the structures most sensitive to change. This study included 203 subjects - 70 bvFTD and 133 CNCs - with a total of 482 timepoints from the Frontotemporal Lobar Degeneration Neuroimaging Initiative.

**Results:** Deformation based morphometry (DBM) revealed significant atrophy in the frontal lobes, insula, medial and anterior temporal regions bilaterally in bvFTD subjects compared to controls with outstanding subcortical involvement. We provide detailed information on regional changes per year. In both cross-sectional analysis and over a one-year follow-up period, ventricle expansion was the most prominent differentiator of bvFTD from controls and a sensitive marker of disease progression.

**Conclusions:** Automated measurement of ventricular expansion is a sensitive and reliable marker of disease progression in bvFTD to be used in clinical trials for potential disease modifying drugs, as well as possibly to implement in clinical practice. Ventricular expansion measured with DBM provides the lowest published estimated sample size for clinical trial design to detect significant differences over one and two years.

## 1. Introduction

In an effort to address the need for improved diagnostic biomarkers for the behavioural variant frontotemporal dementia (bvFTD), several studies have demonstrated the potential value of morphometric MRI analysis for diagnostic purposes (McCarthy, Collins et al. 2018). Here, we performed a deformation-based morphometry (DBM) study of longitudinal MRI changes in bvFTD.

Unlike voxel-based morphometry (VBM) where sometimes erroneous tissue classification can lead to incorrect measurement of gray matter volume, DBM does not depend on the automated segmentation into gray matter, white matter and CSF (Ashburner, Hutton et al. 1998, Ashburner and Friston 2000, Chung, Worsley et al. 2001). Instead, it uses image contrast directly as an explicit representation of these distributions. The improvements on nonlinear image registration algorithms allow for matching the images locally based on similarities in contrast and intensities, making DBM more sensitive than VBM for subtle differences. In addition, the image processing tools used in this study have been designed to process data from multi-site studies to handle biases due to multi-site scanning and have been applied successfully to a number of multi-site projects (Zeighami, Ulla et al. 2015, Boucetta, Salimi et al. 2016, Dadar, Fonov et al. 2018, Dadar, Maranzano et al. 2018).

In this study, our objectives were: to precisely quantify the magnitude and pattern of volume change in bvFTD as compared to cognitively normal controls (CNCs) using DBM that rely on robust registration methods, to compare the progression of atrophy between these two cohorts over one year follow-up and to identify the structures most sensitive to change and generate sample size estimates for these regions of interest (ROIs) for the design of future therapeutic trials in bvFTD.

## 2. Materials and Methods

### 2.1 Participants

The frontotemporal lobar degeneration neuroimaging initiative (FTLDNI) was funded through the National Institute of Aging and started in 2010. The primary goals of FTLDNI are to identify neuroimaging modalities and methods of analysis for tracking frontotemporal lobar degeneration (FTLD) and to assess the value of imaging versus other biomarkers in diagnostic roles. The project is the result of collaborative efforts at three sites in North America. For up-to-date information on participation and protocol, please visit: http://4rtni-ftldni.ini.usc.edu/

Data was accessed and downloaded through the LONI platform in August 2018. We included bvFTD patients and CNCs from the FTLDNI database who had T1-weighted (T1w) MRI scans matching with each clinical visit. The inclusion criteria for bvFTD patients was diagnosis of possible or probable bvFTD according to the frontotemporal dementia (FTD) consortium criteria (Rascovsky, Hodges et al. 2011). All subjects included provided informed consent and the protocol was approved by the institution review board at all sites.

### 2.2. Clinical assessment

All subjects were assessed in periodic visits (every 6 months) for clinical characteristics (motor, non-motor and neuropsychological performance) by site investigators. Neuropsychological assessment included Mini Mental State Examination (MMSE), Montreal Cognitive Assessment (MOCA), Clinical Dementia Rating Scale (CDR) global score, FTLD Clinical Dementia Rating Scale Sum of Boxes (CDR-SOB), Clinical Global Impression (CGI), California Verbal Learning Test (CVLT), Modified Trials (MTT), Digit span forward and backward, Verbal Fluency, Neuropsychiatry Inventory, Functional Activities Questionnaire (FAQ), Boston Naming Test (BNT), Behavioral Activation Scale (BAS), Behavioral Inhibition Scale (BIS), Schwab and England Activities of Daily Living Scale (SEADL).

### 2.3. Clinical data analysis

All statistical analyses were conducted using MATLAB (version R2018b). Two-sample t-Tests were conducted to compare demographic and clinical variables at baseline. Categorical variables (e.g., sex) were analysed using chi-square analyses. Results are expressed as mean ± standard deviation and median [interquartile range]. A two-sided p-value of <0.05 was considered statistically significant.

### 2.4. Neuroimaging

#### 2.4.1. Image Acquisition

3.0T MRIs were acquired on three sites. The acquisition parameters for each site are summarized in Table 1. In all sites, volumetric MPRAGE sequence was used to acquire T1w images of the entire brain.

**Table 1.**
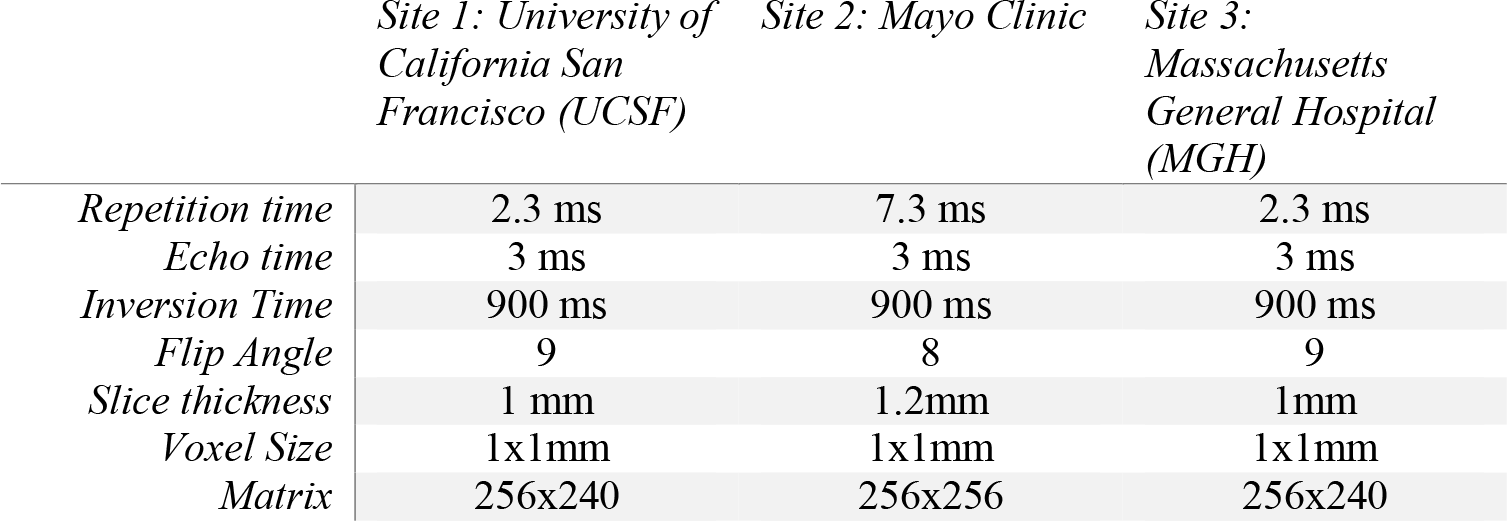
Structural T1-weighted image acquisition protocols by center.

#### 2.4.2. Pre-Processing

All T1w scans for each subject were pre-processed through our local longitudinal pipeline (Aubert-Broche, Fonov et al. 2013). Image denoising (Coupe, Yger et al. 2008), intensity non-uniformity correction (Sled, Zijdenbos et al. 1998), and image intensity normalization into range (0−100) using histogram matching were performed. For each subject, each native T1w volume from each timepoint was linearly registered together to form a subject-specific template, aligned with the ICBM152 template (Collins, Neelin et al. 1994, A.C. Evans 1997). Each T1w volume was then non-linearly registered to the ICBM152 template using ANTs diffeomorphic registration pipeline (Avants, Epstein et al. 2008). The quality of the registrations was visually assessed and cases that did not pass this quality control were discarded (n=16 scans).

#### 2.4.3. Deformation based morphometry

DBM analysis was performed using MNI MINC tools. The principle of DBM is to warp each individual scan to a common template by high-dimensional deformation, where shape differences between the two images (i.e., the subject’s T1w and the template) are encoded in the deformations. The local deformation obtained from the non-linear transformations was used as a measure of tissue expansion or atrophy by estimating the determinant of the Jacobian for each transform. Local contractions can be interpreted as shrinkage of tissue (atrophy) and local expansions as growing are often related to ventricular or sulci enlargement. DBM was used to assess both voxel-wise and atlas-based cross-sectional group related volumetric differences. In addition, we evaluated longitudinal change and correlation with disease staging scores.

#### 2.4.4. Voxel-Wise analysis

A voxel-wise mixed effects model analysis was performed to assess the pattern of volumetric change over time according to diagnosis. The following model was used:

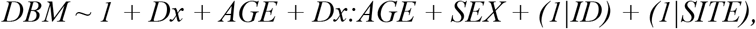

where DBM values are 193*229*193 Jacobian matrices for different subject timepoints. Dx is a categorical variable for bvFTD versus CNCs. Sex is also a categorical variable. ID and Site are categorical random effects. The term Dx:AGE denotes the interaction between diagnostic group and age. The Jacobian determinant from the DBM analysis (as a proxy of local atrophy) was the dependent variable and age, sex, and diagnosis were the independent fixed variables. The model also included an interaction term between diagnosis and age in order to assess the additional impact of age on bvFTD compared to CNCs. The resulting maps were corrected for multiple comparisons using False Discovery Rate (FDR) (Genovese, Lazar et al. 2002), thresholded at ≤0.05 to identify regions associated with differences between bvFTD and CNCs.

#### 2.4.5. Atlas-based analysis

In order to provide regional information on cerebral changes, an atlas-based analysis was also used to determine the mean volume difference of all cortical and subcortical structures. These structures were identified by manually defined labels based on the Mindboogle-101 labelling protocol registered to the ICBM152 template (Klein and Tourville 2012). For each structure, the mean and standard deviation of the Jacobian values (as a proxy for volume) for each cohort were determined. Then, a two-sample t-test for equal means was performed, comparing the two groups (corrected for age and sex).

The magnitude of atrophy (or expansion) per year was assessed performing multiple linear regressions on the average DBM Jacobian values per region for each group by age for subjects that had more than 1 visit (N=160; 53 bvFTD and 107 CNCs). The same procedure was implemented with the functional and cognitive scores to obtain the mean annual change and sample size estimates for each cohort. In order to determine if the rate of atrophy varied according to disease stage, bvFTD subjects were divided into tertiles according CDR-SOB;≤ 4.5, 4.5-8 and ≥8. Kruskal-Wallis one-way analysis of variance was performed in order to compare the annual change in DBM Jacobian values between the 3 groups. A two-sample t-test was used in order to compare the annual change in ventricular size between CNCs and the mildest bvFTD subjects (as defined previously; FTLD-CDR <4.5).

Finally, using the differences in the mean annual rate of atrophy/enlargement between the two groups in each region, the sample sizes (per arm) necessary to detect a 25% reduction in yearly change due to disease were estimated for use in potential clinical trials (80% power and 5% 1-tailed significance). All estimates were multiplied by 1.20 to account for 20% expected attrition. Annual change in DBM Jacobian values was also correlated with the annual change in CDR-SOB for the structures with sample sizes smaller than 300.

## 3. Results

### 3.1. Demographics

We included 203 subjects; 70 bvFTD and 133 CNCs with a total of 482 timepoints. Table 2 shows the demographic and cognitive testing performances in bvFTD and CNCs. There was no statistically significant difference in mean age between bvFTD patients vs CNCs (62.1±6.5 and 64.1±7.5 years respectively, p=0.06). There was a higher proportion of men in bvFTD patients than CNCs (67% vs 42%, p=0.001). The number of follow-up visits was comparable in the two cohorts, albeit a slightly longer follow-up time in controls vs bvFTD patients (1.15 [0.6-3] vs 1.0 [0.3-1.1] years respectively, p<0.001). As expected, bvFTD subjects showed greater cognitive and functional impairment. Significant differences were found between the two cohorts in MMSE, MOCA, CGI, language and behavior CDR, and both global score CDR and CDR-SOB (p<0.001). Complete neuropsychological test results for bvFTD and CNCs can be found in Table S1 in the Supplementary Material.

**Table 2.**
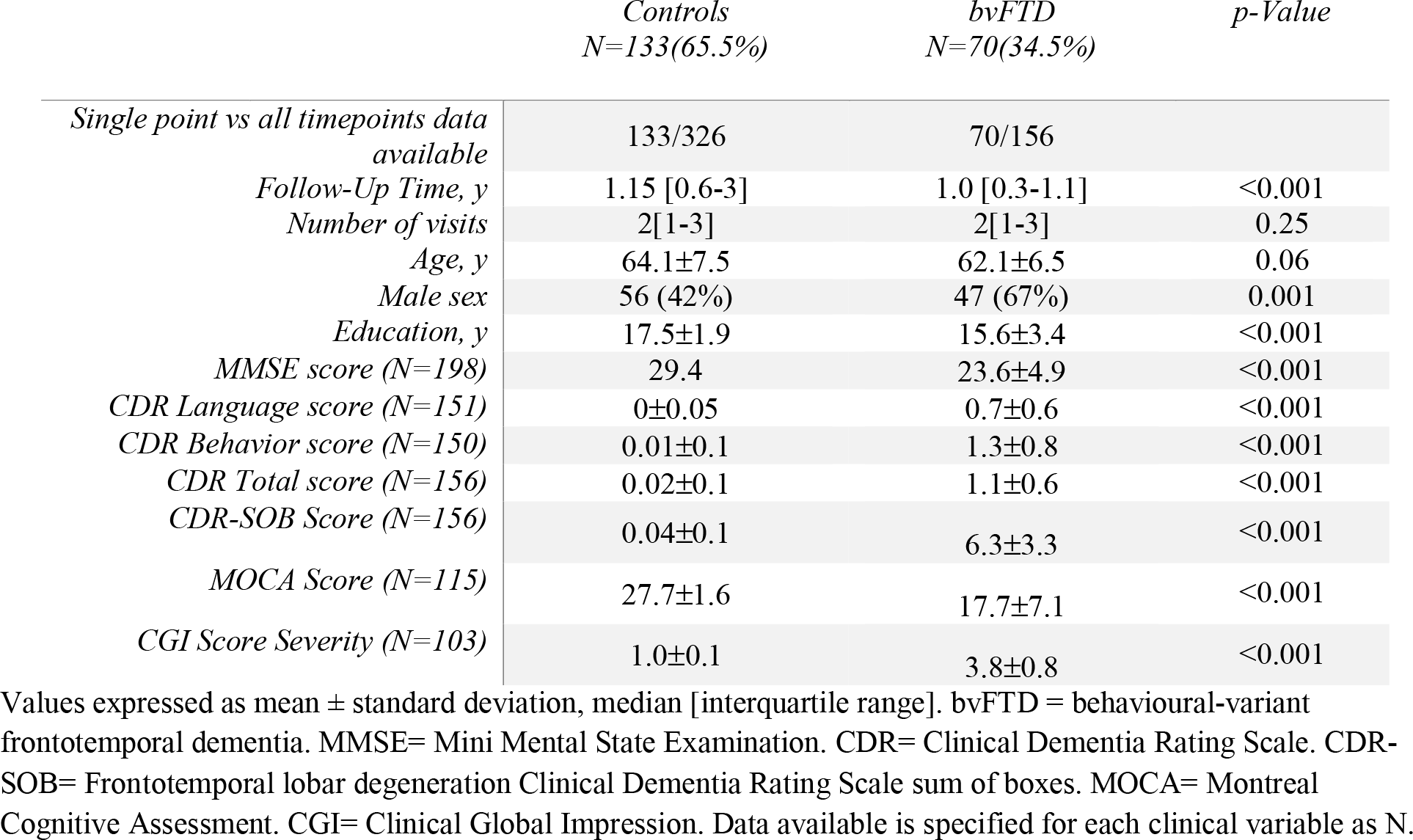
Baseline demographic and clinical characteristics in bvFTD and healthy controls

|  | *Controls*<br>*N=133(65.5%)* | *bvFTD*<br>*N=70(34.5%)* | *p-Value* |
| --- | --- | --- | --- |
| *Single point vs all timepoints data available* | 133/326 | 70/156 |  |
| *Follow-Up Time, y* | 1.15 [0.6-3] | 1.0 [0.3-1.1] | <0.001 |
| *Number of visits* | 2[1-3] | 2[1-3] | 0.25 |
| *Age, y* | 64.1±7.5 | 62.1±6.5 | 0.06 |
| *Male sex* | 56 (42%) | 47 (67%) | 0.001 |
| *Education, y* | 17.5±1.9 | 15.6±3.4 | <0.001 |
| *MMSE score (N=198)* | 29.4 | 23.6±4.9 | <0.001 |
| *CDR Language score (N=151)* | 0±0.05 | 0.7±0.6 | <0.001 |
| *CDR Behavior score (N=150)* | 0.01±0.1 | 1.3±0.8 | <0.001 |
| *CDR Total score (N=156)* | 0.02±0.1 | 1.1±0.6 | <0.001 |
| *CDR-SOB Score (N=156)* | 0.04±0.1 | 6.3±3.3 | <0.001 |
| *MOCA Score (N=115)* | 27.7±1.6 | 17.7±7.1 | <0.001 |
| *CGI Score Severity (N=103)* | 1.0±0.1 | 3.8±0.8 | <0.001 |

Values expressed as mean ± standard deviation, median [interquartile range]. bvFTD = behavioural-variant frontotemporal dementia. MMSE= Mini Mental State Examination. CDR= Clinical Dementia Rating Scale. CDR-SOB= Frontotemporal lobar degeneration Clinical Dementia Rating Scale sum of boxes. MOCA= Montreal Cognitive Assessment. CGI= Clinical Global Impression. Data available is specified for each clinical variable as N.

### 3.2. Voxel-Wise DBM analysis

#### 3.2.1. Cross-sectional

Figure 1 shows the statistically significant differences in local volume differences between bvFTD and CNCs after FDR correction. Greater gray and white matter atrophy are evident in the medial and inferior lateral portions of the frontal lobes as well as dorsolateral prefrontal cortex, insula, basal ganglia, medial and anterior temporal regions bilaterally and regions of brainstem and cerebellum in bvFTD. A corresponding volume increase is shown in the ventricles and sulci, being more evident in frontal horns and lateral sulcus. Restricted bilateral involvement of parietal cortex was also found, though with a lesser degree of atrophy.

**Figure 1.**
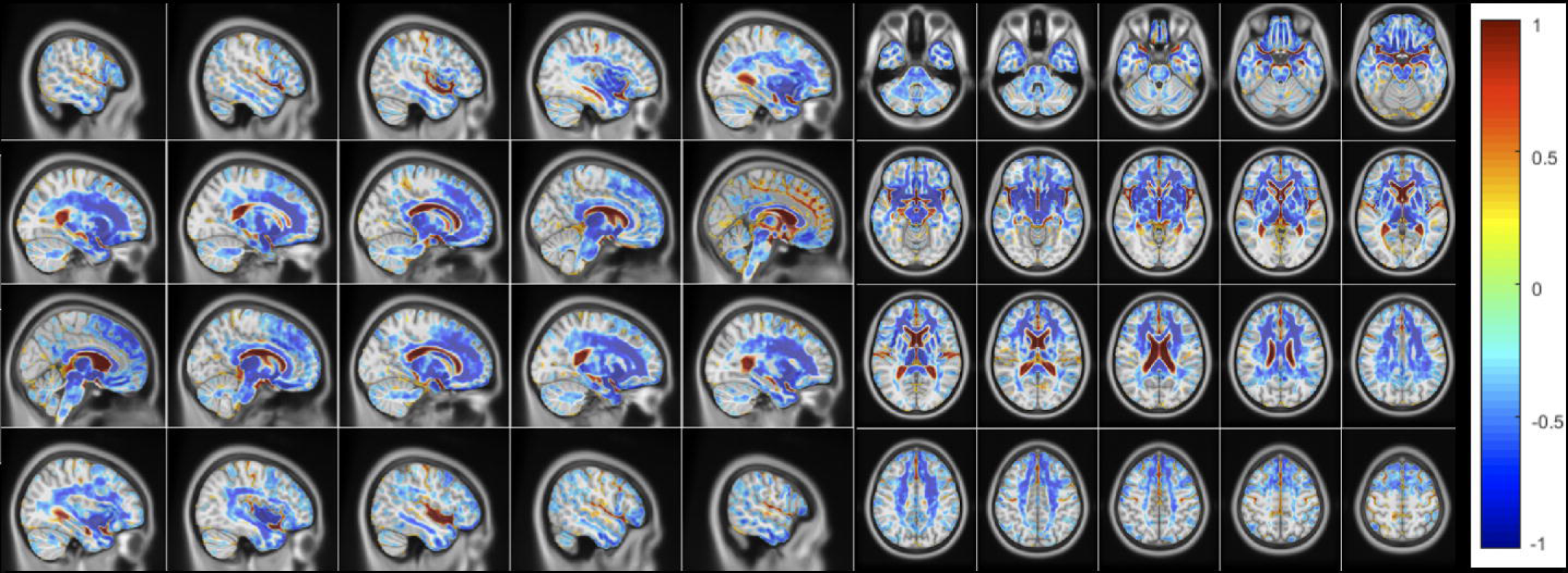
Voxel-wise DBM Jacobian beta maps indicating significant differences between Controls and bvFTD (FDR corrected p-value<0.05). Model: DBM ~ 1 + Dx + AGE + Dx:AGE + Sex + (1|ID) + (1|SITE). The figures show the significant beta values obtained for the categorical variable DX (bvFTD vs Controls). Warmer colors indicate regions with larger DBM values (i.e. ventricle enlargement), and colder colors indicate lower DBM values (i.e. smaller regions) in bvFTD compared to Controls.

#### 3.2.2 Longitudinal

Both bvFTD and CNCs showed predominant medial volume loss due to ageing affecting predominantly white matter (Figure 2). When factoring this age-related volume loss, over a brief follow up period (median 1 year), the enlargement of the frontal horns of the ventricles was the most significant differentiator between bvFTD and CNCs (Figure 3). However, significant atrophy, greater than expected for age, was also found in small areas across the cingulum, callosum and medial frontal cortex in the bvFTD group.

**Figure 2.**
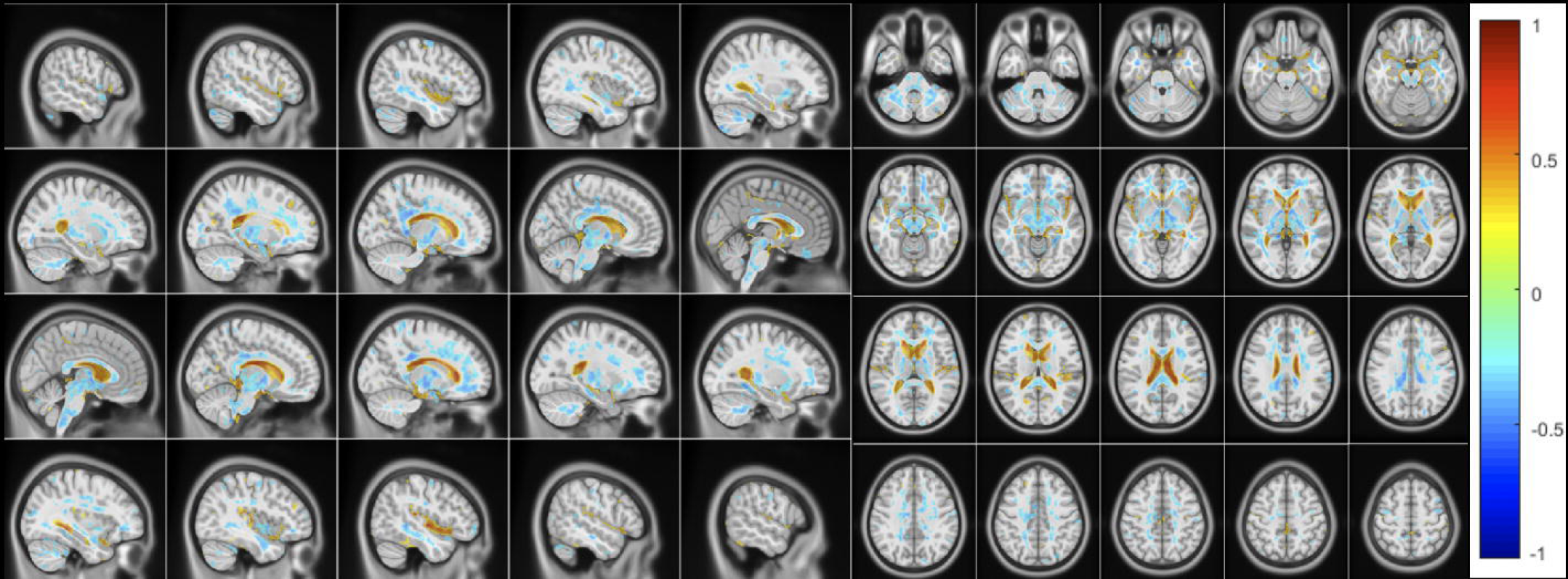
Voxel-wise DBM Jacobian beta maps indicating significant associations with age for the Control cohort only (FDR corrected p-value<0.05). Model: DBM ~ 1 + Dx + AGE + Dx:AGE + SEX + (1|ID) + (1|SITE). The figures show the significant beta values obtained for the continuous variable AGE. Warmer colors indicate regions with larger DBM Jacobian values (i.e. ventricle enlargement), and colder colors indicate smaller DBM Jacobian values (i.e. tissue atrophy) associated with ageing.

**Figure 3.**
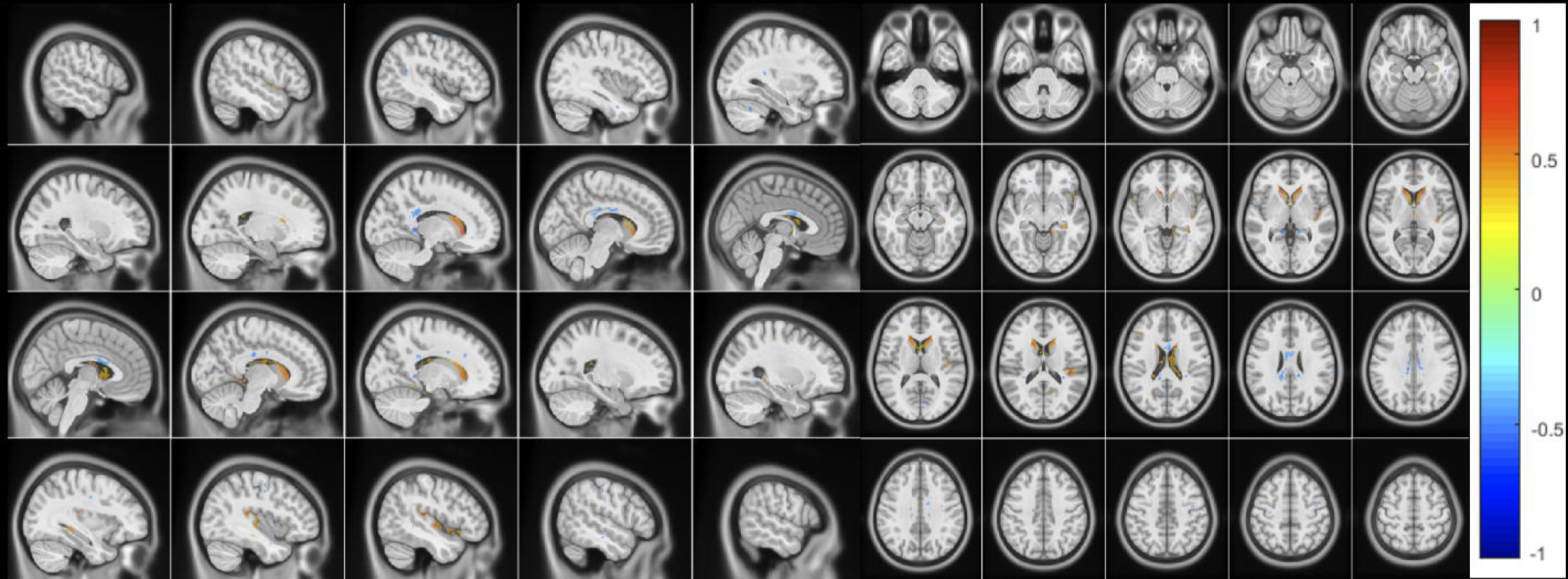
Voxel-wise DBM beta maps indicating additional associations with age for the bvFTD cohort (FDR corrected p-value<0.05). Model: DBM ~ 1 + DX + AGE + DX:AGE + GENDER + (1|ID) + (1/SITE). The figures show the significant beta values obtained for the interaction variable DX:AGE, and indicate the regions of change due to the disease, over and above what is expected for age. Warmer colors indicate regions with larger DBM Jacobian values (i.e. ventricle enlargement), and colder colors indicate smaller DBM Jacobian values (i.e. tissue atrophy) associated with additional effects of ageing in the bvFTD cohort.

### 3.3. Atlas based DBM analysis

#### 3.3.1. Cross-sectional

Assessing the baseline differences per anatomically defined regions between the two cohorts, significant differences were found in a large number of regions with an antero-posterior gradient. The greatest magnitude of change found was the enlargement of lateral and third ventricles (p=<0.001, t>10) and atrophy of the thalamus (p=<0.001, t= −10.62), followed by significant atrophy of the amygdala, lateral orbitofrontal cortex and putamen (p=<0.001, t= −9.76, t= −9,69, t= −9.5 respectively). Table 3 shows the mean DBM Jacobian values, p-values and t-values for the structures with the largest (t-values ≥ 5) differences between bvFTD and CNCs after FDR correction. The complete table with all the statistically significant structures can be found in Supplementary table S2.

**Table 3.**
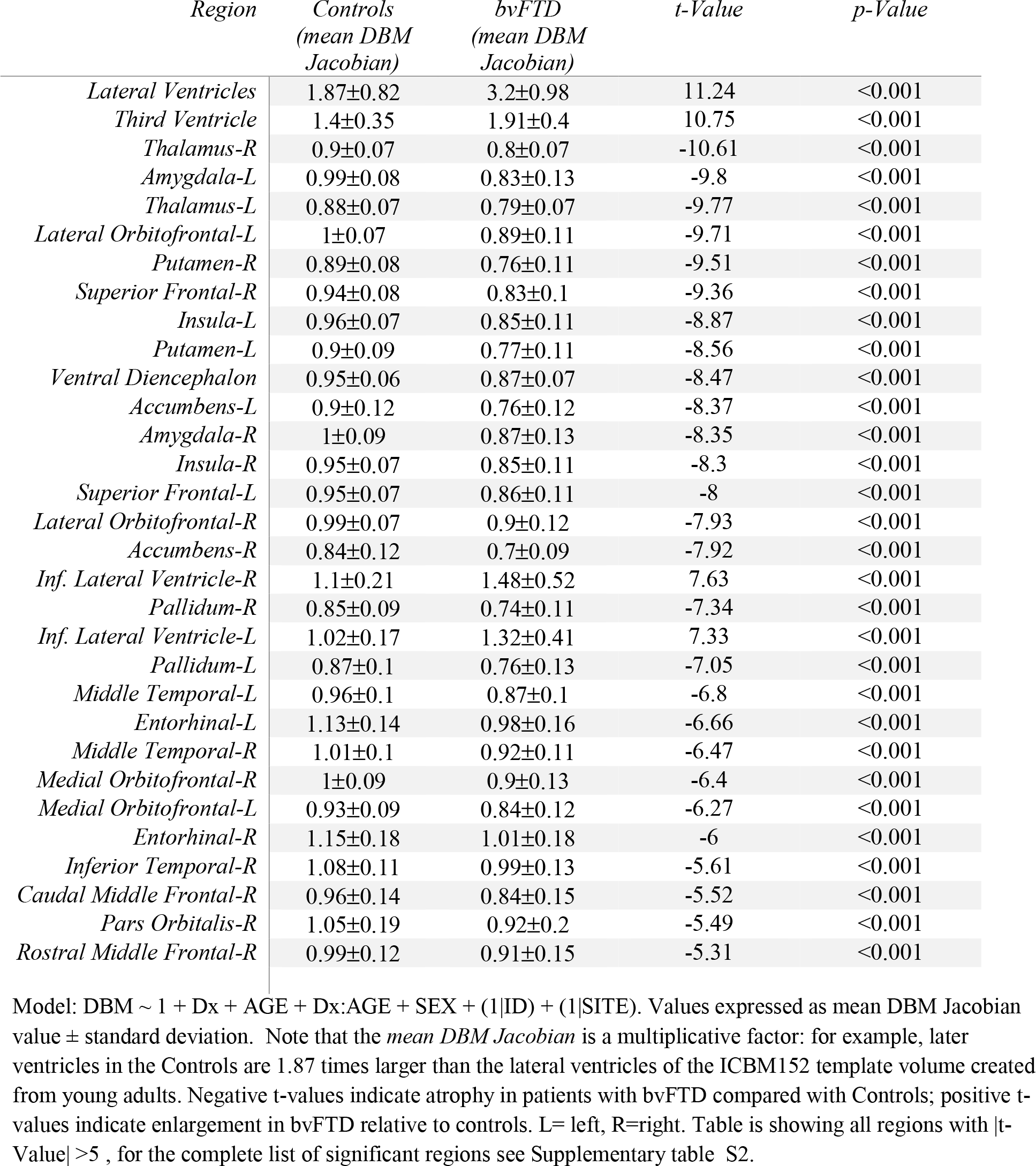
Regions with statistically significant cross-sectional differences at baseline between bvFTD and Controls, sorted by t-Value magnitude.

#### 3.3.2 Annual change and sample size estimation

Table 4 lists the structures that showed significant differences in progression of atrophy/enlargement in the 1-year follow up; the mean annual changes of DBM per region for each cohort and the corresponding p-values. Many cortical and subcortical structures showed significant progression in atrophy (i.e., reduction in DBM Jacobian value) in bvFTD compared to CNCs. However, the greatest change in one year was associated to the enlargement of the ventricles with a mean annual change 4 to 10 times greater for bvFTD than CNCs for lateral ventricles, third ventricle and inferior lateral ventricles (p<0.001). As mentioned, statistically significant differences in the annual change between the groups were also found in several cortical and subcortical structures such as posterior cingulate, diencephalon, putamen and thalamus (p<0.001).

**Table 4.**
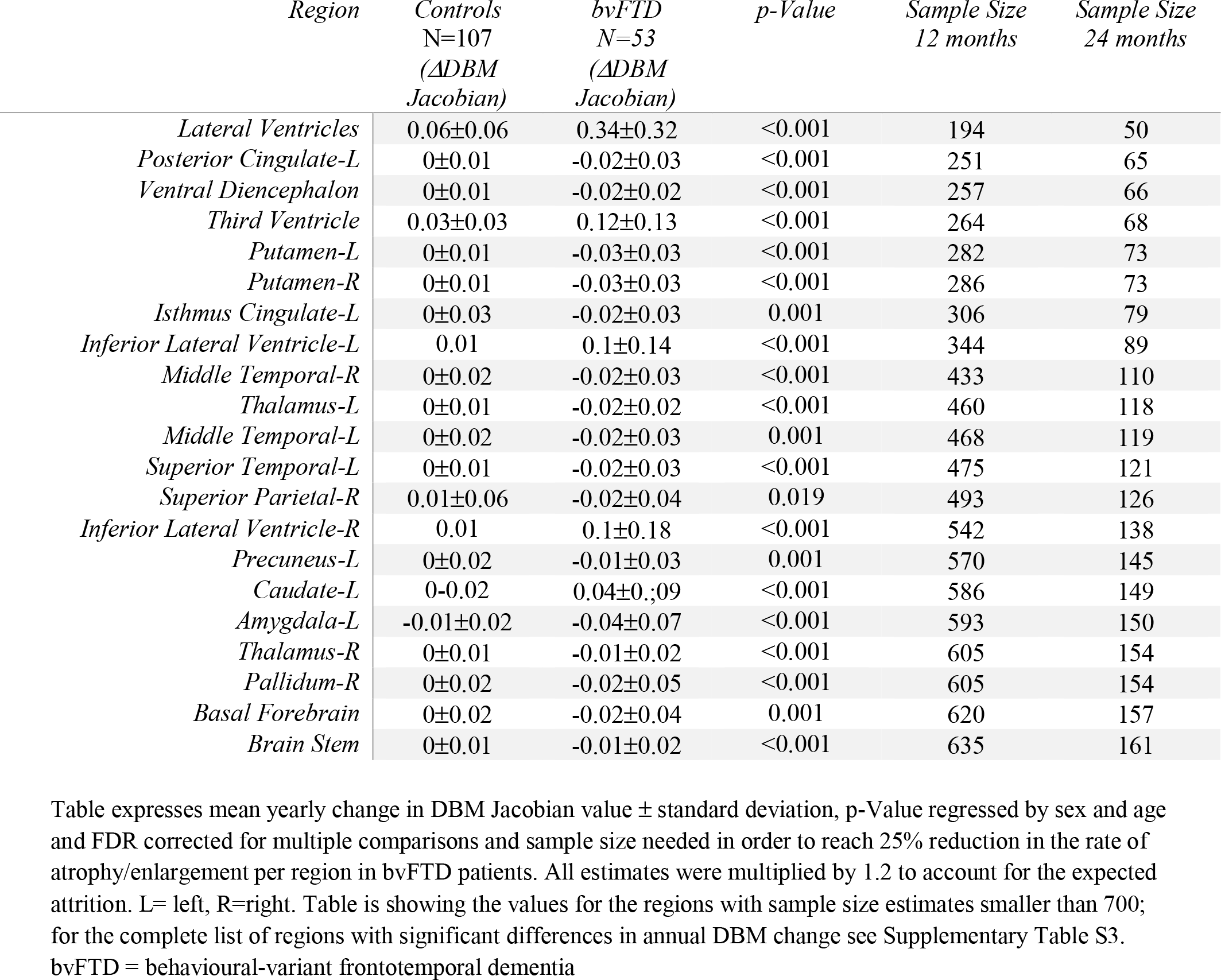
Regions with statistically significant differences in progression of atrophy/enlargement in 1 year follow up between bvFTD and Controls, classified by sample size estimates

Table 4 also compares the sample size estimates per arm to detect 25% reduction in the rate of progression of atrophy/enlargement with 80% power and 0.05 level of significance in a hypothetical 12 and 24-month follow-up, 1:1 parallel group clinical trial and expected attrition of 20%. The sample size estimated to detect a 25% reduction in the enlargement of the ventricles in one year is 194 patients per arm for the lateral ventricles; followed by 251 for the posterior cingulate gyrus, 257 for diencephalon and 264 for the third ventricle. For a 24-months trial the estimated sample sizes to detect a 25% reduction in the rate of progression of atrophy/enlargement are 50, 65, 66 and 68 respectively for the same structures.

#### 3.3.3 Ventricular annual change and disease severity

Figure 4 panel A shows the change in ventricular volume (VV) per year according to CDR-SOB groups. Using Kruskal-Wallis one-way analysis of variance, there were no significant differences in annual change in ventricular expansion between the three bvFTD groups (i.e. ΔDBM Jacobian, p= 0.11). However, all three bvFTD groups showed statistically significant differences compared to the CNCs group (p<0.001). Figure 4 panel B plots the comparison in between CNCs and bvFTD subjects with CDR-SOB ≤ 4.5 (i.e, milder forms of the disease), confirming that early stage bvFTD patients have a ΔDBM Jacobian significantly higher than controls (p<0.001).

**Figure 4.**
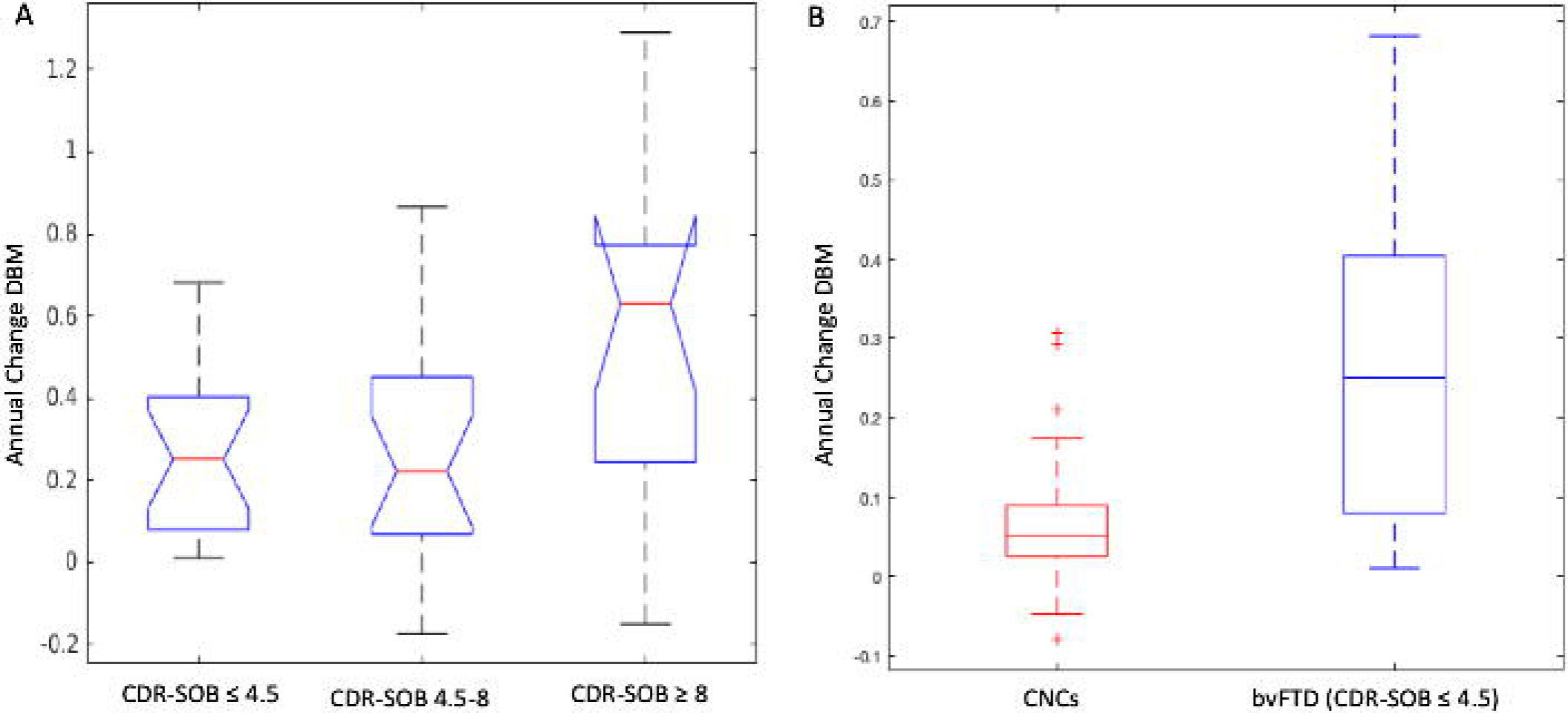
**Panel A**. Boxplots comparing the annual change in DBM Jacobian for the lateral ventricles of the three bvFTD groups (CDR-SOB ≤ 4.5, CDR-SOB 4.5-8 and CDR-SOB ≥ 8). **Panel B.** Boxplots comparing the annual change in DBM Jacobian for the lateral ventricles between controls and bvFTD subjects with CDR-SOB ≤ 4.5. CDR-SOB: clinical dementia rating score sum of boxes; DBM: deformation-based morphometry. CNCs: cognitively normal controls. bvFTD: behavioral variant frontotemporal dementia

### 3.4 Clinical severity analyses

Significant differences were found between bvFTD and CNCs for the mean annual change in CDR-SOB, MMSE, MOCA and CGI scores corrected by age and gender (Supplementary table S4). The sample sizes estimated to detect a 25% improvement on each of these clinical measures is 204, 552, 310 and 720 individuals per arm respectively (80% power, 0.05 level of significance and 20% expected attrition).

Figure 5 shows the correlation between annual DBM Jacobian change and annual change in CDR-SOB in bvFTD subjects for selected regions (with significant differences from CNCs and smaller sample sizes). Significant correlations were found for lateral ventricles, third ventricle and diencephalon, with the strongest correlation found for lateral ventricles (p=0.005, r=0.37).

**Figure 5.**
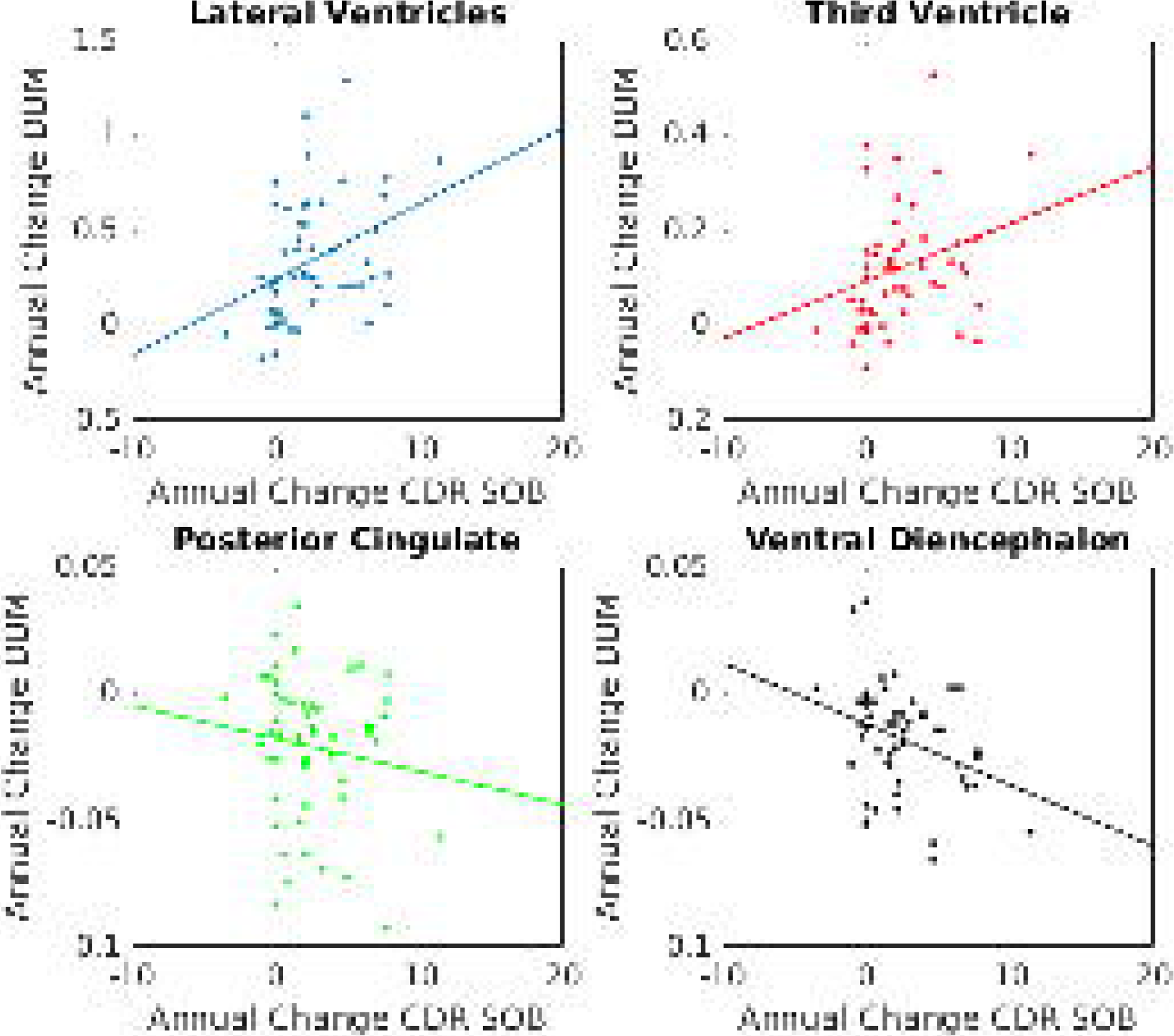
Plots representing annual DBM change vs annual change in CDR-SOB sum of for lateral ventricles (p=0.005, r=0.37), third ventricle (p=0.04; r=0.28), posterior cingulate (p=ns, r=−0.14) and diencephalon (p=0.01, r=−0.34). CDR-SOB: clinical dementia rating score sum of boxes; DBM: deformation-based morphometry.

## 4. Discussion

The major findings of this study are: (1) subjects with bvFTD at baseline showed the expected atrophy in the frontal lobes (most evident in medial portions), insula, basal ganglia, medial and anterior temporal regions bilaterally both on voxel wise analysis and with anatomically defined ROIs approach; (2) subcortical structures were notably affected in our bvFTD cohort; (3) ventricles and sulci within frontotemporal regions were larger in bvFTD compared to CNC and showed significant enlargement and over a one year period. Ventricular expansion (particularly the lateral ventricles) was the most prominent differentiator of bvFTD from CNCs and thus could be a sensitive marker of disease progression.

From the cross-sectional perspective, the frontal and anterior temporal atrophy observed in bvFTD is consistent with previous post-mortem and voxel-based morphometric studies (Cardenas, Boxer et al. 2007, Whitwell, Boeve et al. 2015, Landin-Romero, Kumfor et al. 2017). Although we did not observe any significant atrophy in occipital lobes in the patients with bvFTD, we did find significant bilateral parietal involvement, though with lesser degree of atrophy compared to frontal and temporal structures. Of particular importance is the involvement of the amygdala and subcortical structures (thalamus, putamen, ventral diencephalon, accumbens area), which in our bvFTD cohort showed the greatest group differences, comparable only to lateral and third ventricle - also an expression of central atrophy - and insular, superior frontal and orbitofrontal cortices.

Over an overall median follow-up time of 1 year, the voxel-wise beta maps showed that the enlargement of the lateral ventricles was the most significant differentiator between bvFTD and CNCs. However, significant progression in atrophy was also found in small areas mainly across the cingulum and medial frontal cortex, consistent with previous longitudinal results with similar follow up period (Brambati, Renda et al. 2007). Unexpected positive values suggesting growth over time were observed for the left caudate (Table 4). This is likely due to a methodological error related to its prominent shrinkage and the corresponding sizeable enlargement of the lateral ventricles, leading to the contamination of the caudate signal with that from the ventricle through partial volume effects. This is a methodological limitation of voxel-based techniques, that has previously been observed in other studies as well (Hua, Leow et al. 2008, Koikkalainen, Lotjonen et al. 2011).

Finally, we found that the ventricles played a remarkable role in discriminating bvFTD from controls and proved to be a sensitive indicator of disease progression. Although this has been suggested in a previous study with fewer bvFTD subjects (Knopman, Jack et al. 2009), our results further confirm these findings and prove the relevance of measuring ventricle enlargement, a finding frequently overlooked in clinical practice. The lateral ventricles appear to be the structure with the most substantial change in DBM Jacobian values per year. Figure 6 shows a visual example of the magnitude of ventricular expansion and caudate atrophy that can occur over 1 year in bvFTD patients. Furthermore, the sample size needed per arm in order to measure 25% decline in the rate of progression of the disease over 12- and 24-months trial is the smallest of all regions examined. Indeed, it is also smaller than the sample size estimated for functional scores and the annual change of the lateral ventricles has shown to correlate significantly with disease severity according CDR-SOB. In addition, the ventricles are a large anatomical area than can be readily measured with volumetric pipelines, and therefore has excellent potential to be integrated as a reliable surrogate marker in clinical trials.

**Figure 6.**
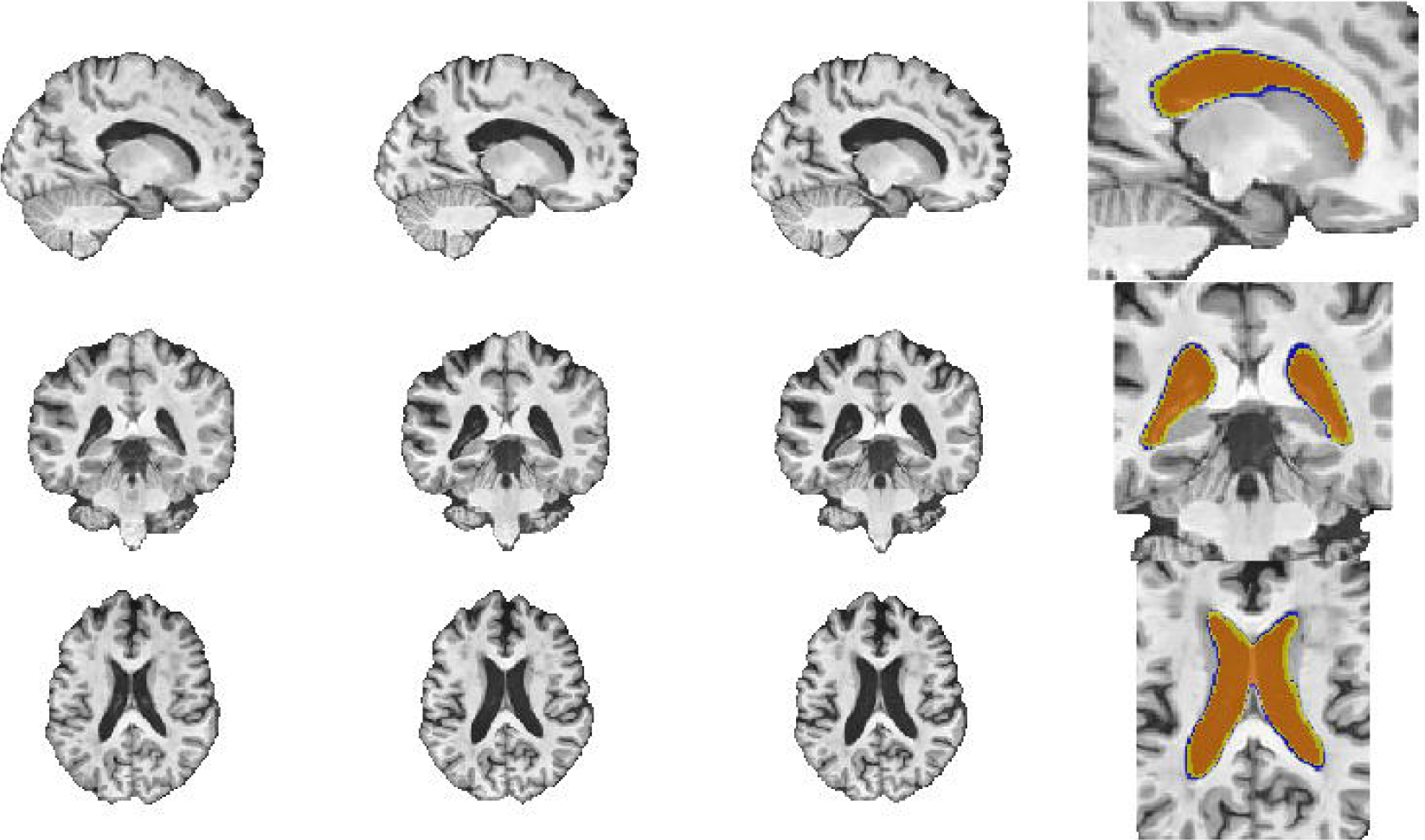
Columns 1-3 correspond to yearly visits 1 to 3 respectively for the same bvFTD subject. Column 4 shows the ventricles overlaid; orange= ventricles at visit 1, green= ventricle enlargement between visit 1 and 2, blue= ventricle enlargement between visit 2 and 3.

Ventricular expansion appears to be closely related to atrophy of subcortical grey matter structures as well as white matter loss. Indeed, it has been reported that dilation of lateral ventricles is preceded by significant atrophy of the basal ganglia, and concurrent to thinning of the corpus callosum (Kril, Macdonald et al. 2005). Though ventricular enlargement is not specific to FTD, it has been found that FTD subjects had higher rates of expansion at all time points compared to Alzheimer’s disease (Whitwell, Jack et al. 2007, Whitwell, Jack et al. 2008, Knopman, Jack et al. 2009). In other studies, bvFTD has shown greater subcortical atrophy at baseline and over time, which is consistent with link between subcortical atrophy and ventricular expansion (Landin-Romero, Kumfor et al. 2017).

According to a previously proposed clinicopathological scheme for staging FTD (Broe, Hodges et al. 2003, Kril and Halliday 2004), our results are in concordance with stage 2, where progression of atrophy is focused in orbital and superior medial frontal cortex along with flattening of the caudate nucleus, and with stage 3 where the hallmark is the involvement of the basal ganglia and ventricles appear considerably dilatated. It has been suggested in previous studies that the rate of brain volume change is not linear throughout the disease (Whitwell, Jack et al. 2008). Furthermore, it has been suggested that trend of progression of atrophy/enlargement might not be the same for different structures. In the present study we assessed the rate of ventricular expansion at different stages. Although bvFTD subjects with CDR-SOB ≥ 8 showed a numerically greater enlargement of the ventricles per year, this difference was not statistically significant. In addition, comparing the rate of progression of mild bvFTD vs CNCs (Figure 4, panel B) also showed that the mentioned potential of VV in discriminating bvFTD from controls could be applicable in the early stages of the disease.

Some studies have shown comparable sample size estimates for clinical trials. Using whole brain and VV annual percent changes as outcome measures for all FTD clinical phenotypes, Knopman et al. estimated 165 bvFTD subjects needed for a 25% effect size and 40% effect size using whole brain volume. Whereas 127 and 51 bvFTD subjects were estimated for 25% and 40% effect size respectively using VV (12-months trial, 80% power, 5% significance and 35% potential attrition rate) (Knopman, Jack et al. 2009). Their study reported 11.2% annual change in VV in a bvFTD cohort (N=34). In comparison, our slightly larger sample size for ventricular annual change could be in relation to different disease severity (our group is slightly older and has an MMSE of 23.6). As mentioned previously, the rate of progression is thought to vary within stages. Other estimates have been made by Binney et al. where the smallest sample size reported is for data-driven ROIs compared to anatomically based ROIs with 409 and 103 subjects per arm for 20 and 40% reduction in rate of decline respectively (12-months trial, 80% power and 5% significance)(Binney, Pankov et al. 2017). Finally, a recent study reports a required sample of 163 bvFTD subjects for 40% effect size and 418 for 25% effect size using left frontal volume as an outcome measure (12-months trial, 80% power, 5% significance and 20% potential attrition rate)(Staffaroni, Ljubenkov et al. 2019); these numbers are more than twice as large are those we estimate for the lateral ventricles. However, when using fractional anisotropy from the corpus callosum, they required only 94 subjects per arm but their estimate was based on diffusion data from a single site. Inter-site variability of diffusion data may increase this number of multi-site trials. Consequently, our analysis shows that using annual change in VV measured by DBM as an outcome measure would require the smallest sample when using morphological data derived from standard T1w MR images. Despite being easier to segment, using larger structures such as whole brain or lobes will require more subjects per arm to detect reduction in the rate of decline. In other words, the use of larger structures may dilute the change signal if atrophy is localized to specific regions, resulting in bigger sample size estimates.

We acknowledge that there are limitations to the present study. First, the short period of follow-up and the lack of disease duration or time from diagnosis, which is not available in the data used for this study. In this regard, we consider that further work with longer follow-up should focus on addressing the possible impact of disease severity on annual change in brain and ventricular volume. Second, DBM is not the most reliable method to assess cortical changes and it is sensitive to partial volume effects. This was demonstrated with the positive values for longitudinal progression in the caudate and is also suspected to affect the effect size of the atrophy in the cortical regions due to sulcal enlargement. Third, compared to cortical thickness, conventional voxel-based techniques are a less sensitive measurement to detect regional gray matter changes related to neurodegeneration at early stages due to partial volume effects (Hutton, Draganski et al. 2009). However, DBM has the advantage of being less sensitive to white matter lesions which are remarkably present in the FTLDNI bvFTD cohort used in this study. Future work will complement these current results with cortical thickness measurements corrected the confounding impact of white matter lesions together with volumetric analysis in order to work out this methodological limitation.

In conclusion, we propose automated measurement of ventricular expansion as a sensitive and reliable marker of disease progression in bvFTD to be used in clinical trials for potential disease modifying drugs, as well as possibly to implement in clinical practice.

## ABBREVIATIONS

bvFTD: behavioural variant of frontotemporal dementia
VBM: voxel-based morphometry
DBM: deformation-based morphometry
CNCs: cognitively normal controls
FTLDNI: frontotemporal lobar degeneration neuroimaging initiative
FTLD: frontotemporal lobar degeneration
T1-w: T1 weighted
FTD: Frontotemporal dementia
MMSE: Mini Mental State Examination
MOCA: Montreal Cognitive Assessment
CDR: Clinical Dementia Rating Scale
CDR-SOB: Frontotemporal lobar degeneration clinical dementia rating scale sum of boxes
CGI: Clinical Global Impression
CVLT: California Verbal Learning Test
MTT: Modified Trials
BNT: Boston Naming test
NPI: Neuropsychiatry Inventory
FAQ: Functional Activities Questionnaire
BAS: Behavioural Activation Scale
BIS: Behavioural Inhibition Scale
SEADL: Schwab and England Activities of Daily Living Scale
MNI-MINC: Montreal Neurological Institute Medical Imaging NetCDF
DX: Diagnosis
FDR: False Discovery Rate
ROIs: Regions of interest
VV: Ventricular volume

## Acknowledgements

We would like to acknowledge funding from the Famille Louise & André Charron. Data collection and sharing for this project was funded by the Frontotemporal Lobar Degeneration Neuroimaging Initiative (National Institutes of Health Grant R01 AG032306). The study is coordinated through the University of California, San Francisco, Memory and Aging Center. FTLDNI data are disseminated by the Laboratory for Neuro Imaging at the University of Southern California.

## Supplementary Material

**Table S1:**
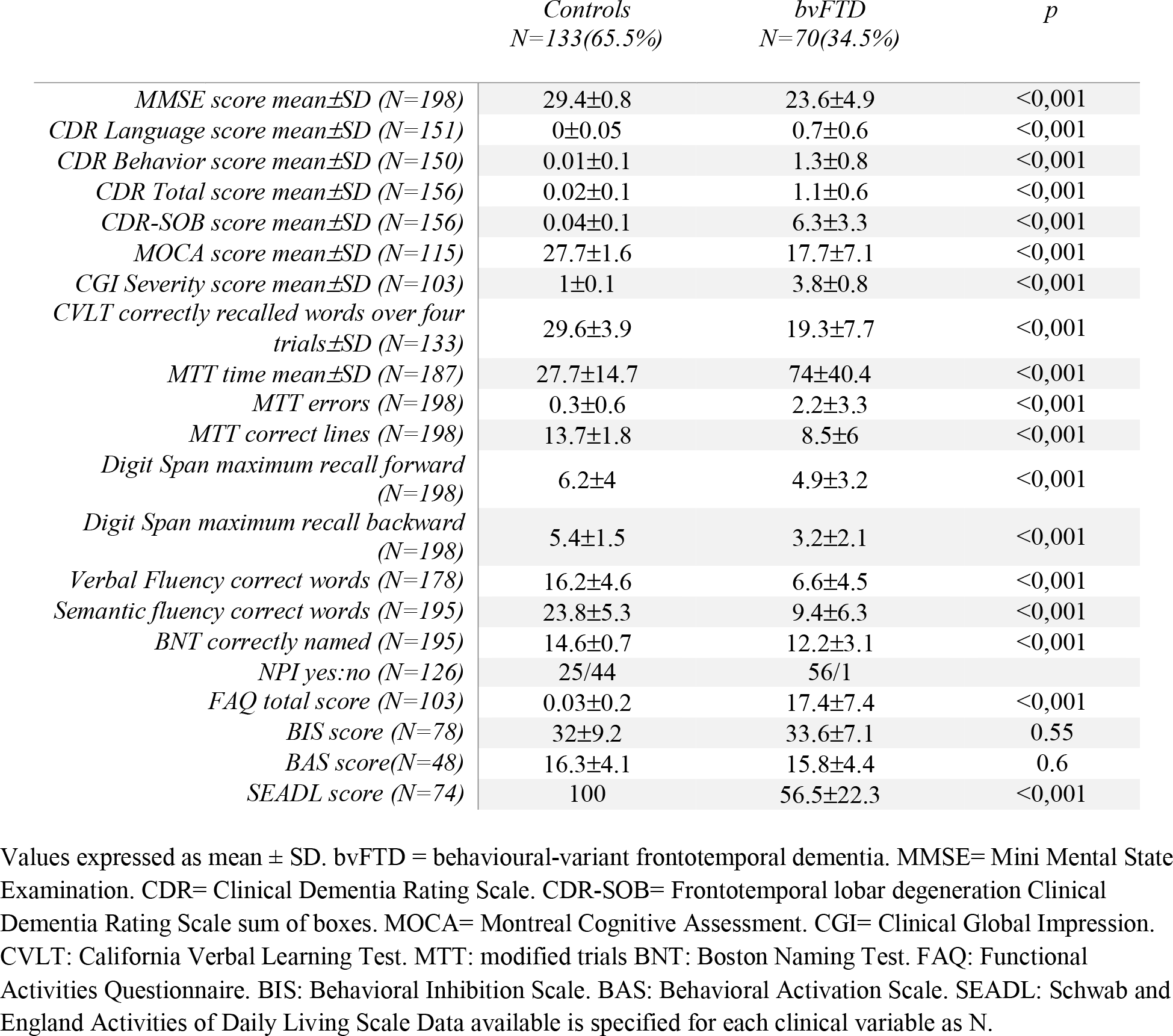
Performance on neuropsychological testing for Controls and bvFTD

**Table S2:**
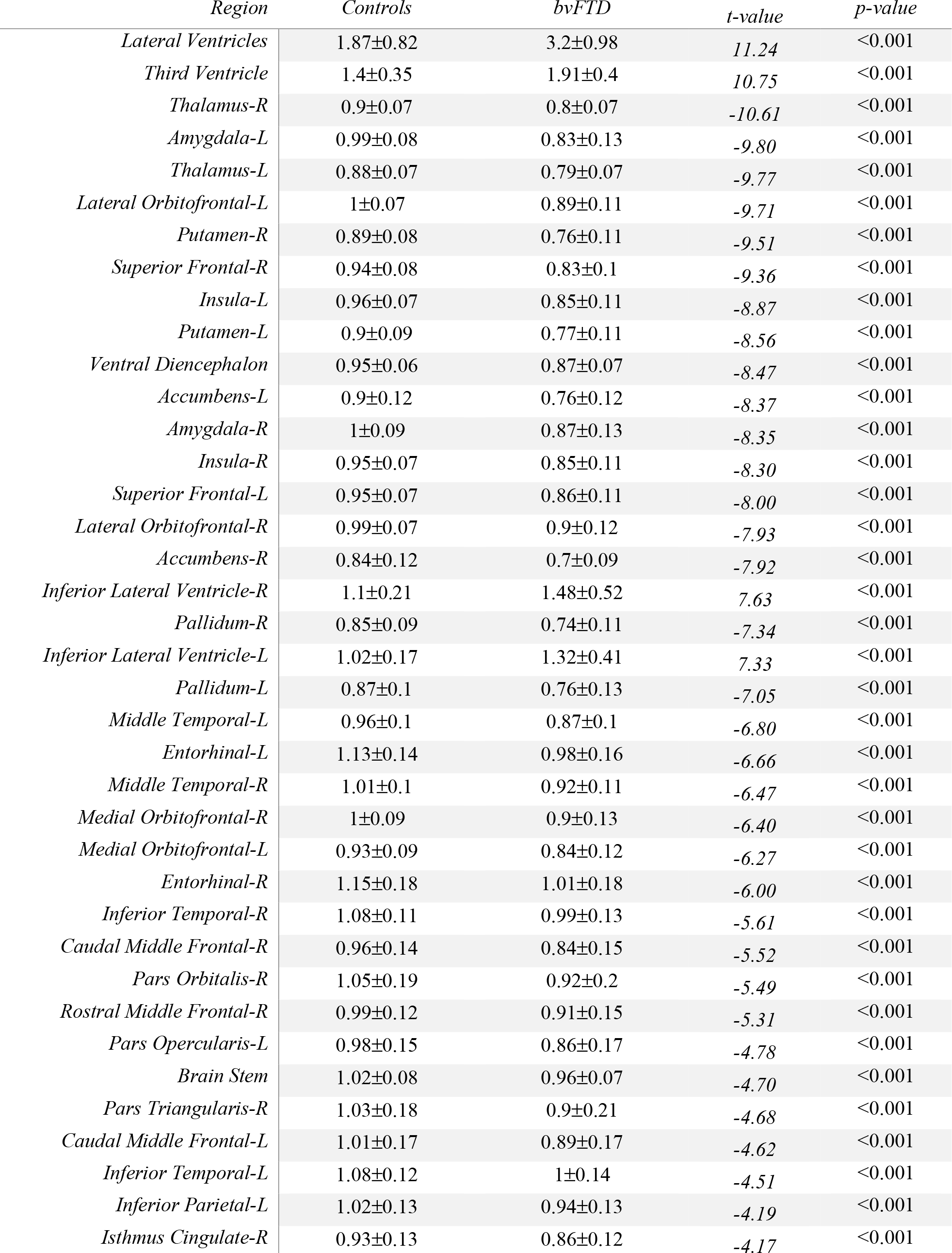
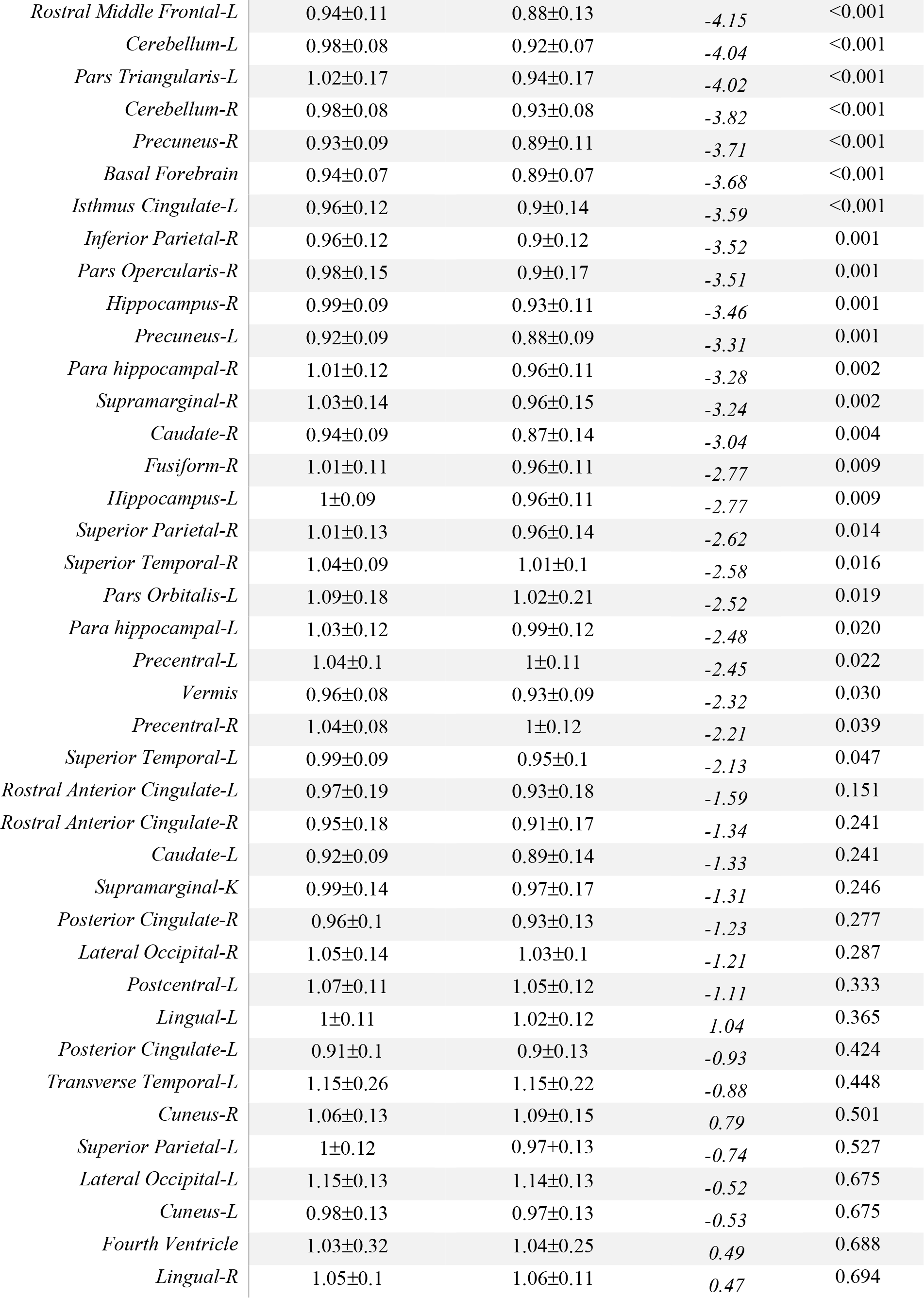
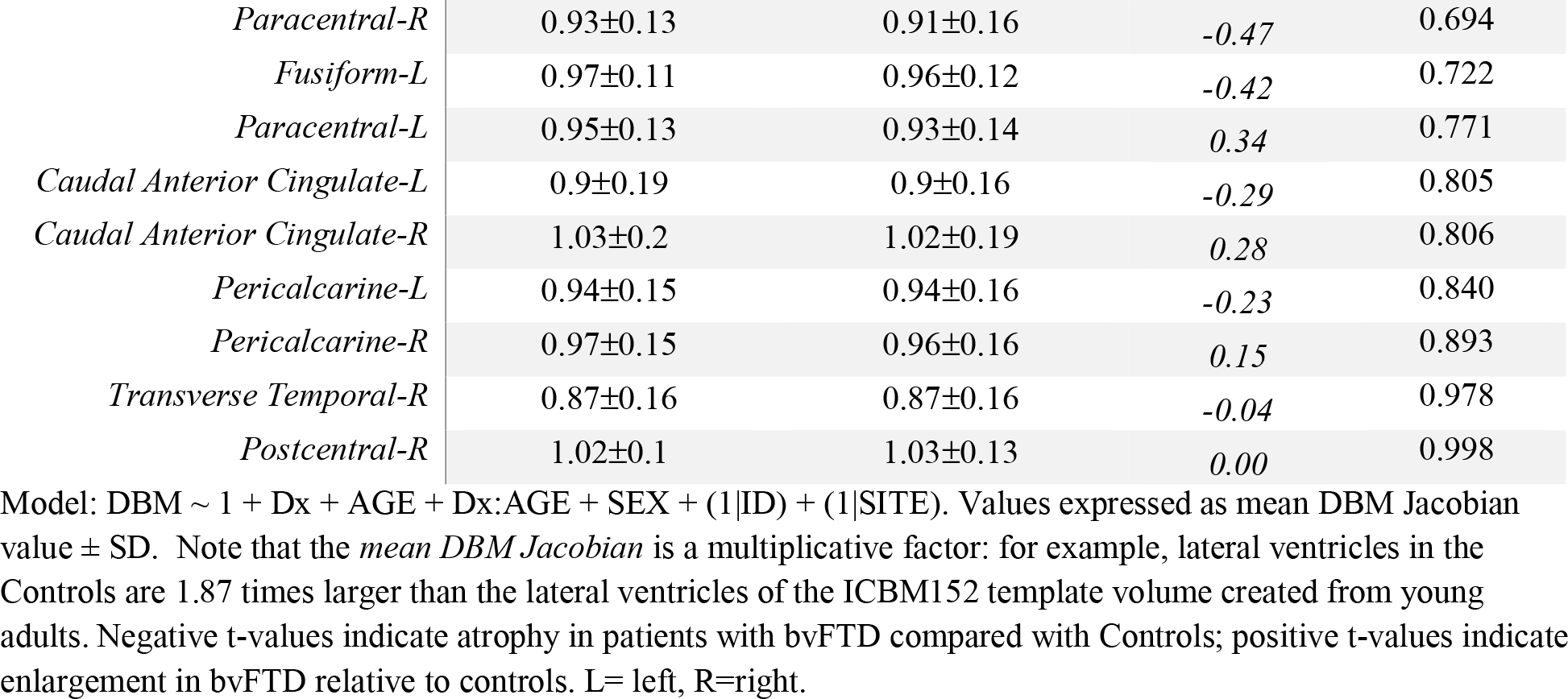
Structures with significant difference between bvFTD and Controls at baseline

**Table S3:**
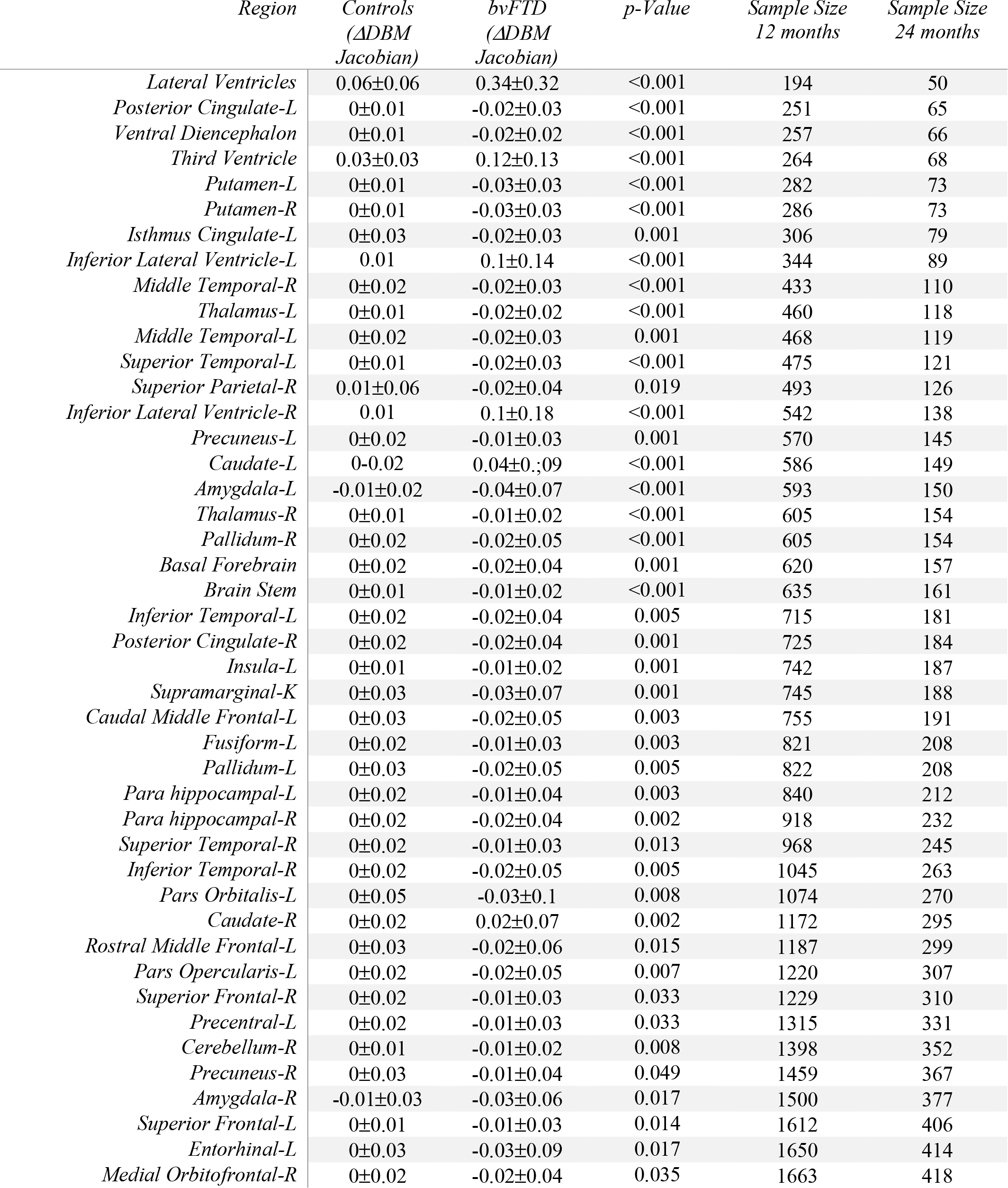
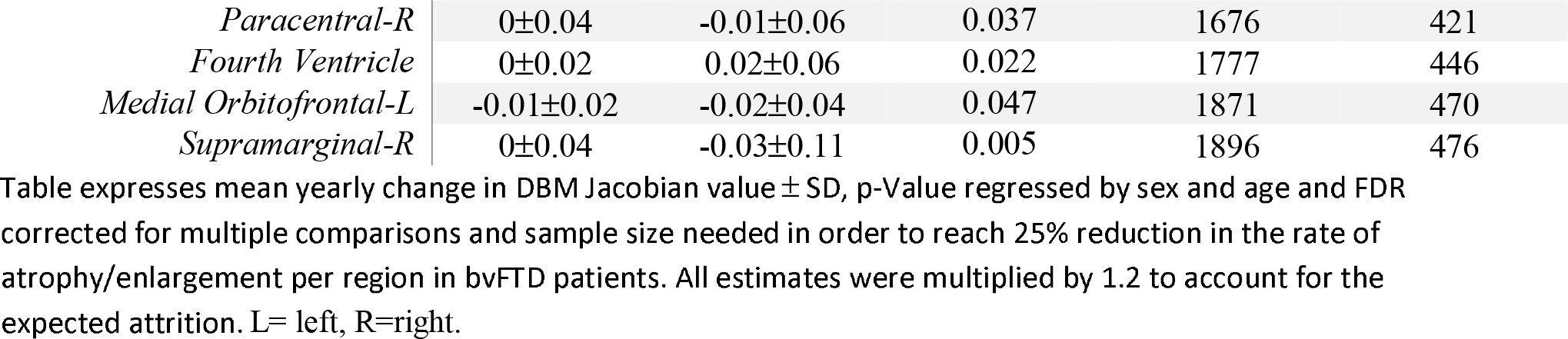
Annual change in DBM per region and sample size estimation both cohorts

**Table S4:**
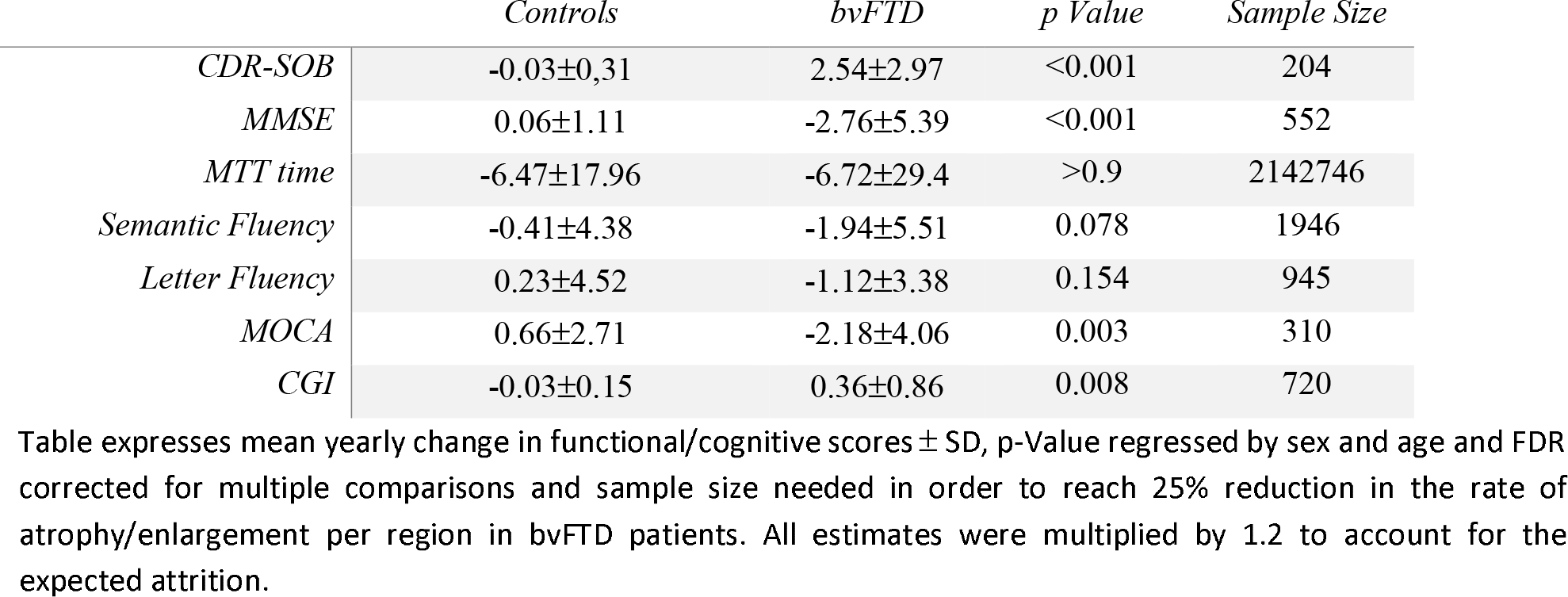
Annual change in clinical scores for bvFTD vs Controls

