## Supplementary material for "DEFORMATION BASED MORPHOMETRY STUDY OF LONGITUDINAL MRI CHANGES IN BEHAVIORAL VARIANT FRONTOTEMPORAL DEMENTIA": Suplemental Material

**Table S1:** Performance on neuropsychological testing for Controls and bvFTD

|  |  | Controls  N=133(65.5%) | bvFTD  N=70(34.5%) | p |
| --- | --- | --- | --- | --- |
| MMSE score mean±SD (N=198) | | 29.4±0.8 | 23.6±4.9 | <0,001 |
| CDR Language score mean±SD (N=151) | | 0±0.05 | 0.7±0.6 | <0,001 |
| CDR Behavior score mean±SD (N=150) | | 0.01±0.1 | 1.3±0.8 | <0,001 |
| CDR Total score mean±SD (N=156) | | 0.02±0.1 | 1.1±0.6 | <0,001 |
| CDR-SOB score mean±SD (N=156) | | 0.04±0.1 | 6.3±3.3 | <0,001 |
| MOCA score mean±SD (N=115) | | 27.7±1.6 | 17.7±7.1 | <0,001 |
| CGI Severity score mean±SD (N=103) | | 1±0.1 | 3.8±0.8 | <0,001 |
| CVLT correctly recalled words over four trials±SD (N=133) | | 29.6±3.9 | 19.3±7.7 | <0,001 |
| MTT time mean±SD (N=187) | | 27.7±14.7 | 74±40.4 | <0,001 |
| MTT errors (N=198) | | 0.3±0.6 | 2.2±3.3 | <0,001 |
| MTT correct lines (N=198) | | 13.7±1.8 | 8.5±6 | <0,001 |
| Digit Span maximum recall forward (N=198) | | 6.2±4 | 4.9±3.2 | <0,001 |
| Digit Span maximum recall backward (N=198) | | 5.4±1.5 | 3.2±2.1 | <0,001 |
| Verbal Fluency correct words (N=178) | | 16.2±4.6 | 6.6±4.5 | <0,001 |
| Semantic fluency correct words (N=195) | | 23.8±5.3 | 9.4±6.3 | <0,001 |
| BNT correctly named (N=195) | | 14.6±0.7 | 12.2±3.1 | <0,001 |
| NPI yes:no (N=126) | | 25/44 | 56/1 |  |
| FAQ total score (N=103) | | 0.03±0.2 | 17.4±7.4 | <0,001 |
| BIS score (N=78) | | 32±9.2 | 33.6±7.1 | 0.55 |
| BAS score(N=48) | | 16.3±4.1 | 15.8±4.4 | 0.6 |
| SEADL score (N=74) | | 100 | 56.5±22.3 | <0,001 |

Values expressed as mean ± SD. bvFTD = behavioural-variant frontotemporal dementia. MMSE= Mini Mental State Examination. CDR= Clinical Dementia Rating Scale. CDR-SOB= Frontotemporal lobar degeneration Clinical Dementia Rating Scale sum of boxes. MOCA= Montreal Cognitive Assessment. CGI= Clinical Global Impression. CVLT: California Verbal Learning Test. MTT: modified trials BNT: Boston Naming Test. FAQ: Functional Activities Questionnaire. BIS: Behavioral Inhibition Scale. BAS: Behavioral Activation Scale. SEADL: Schwab and England Activities of Daily Living Scale Data available is specified for each clinical variable as N.

**Table S2:** Structures with significant difference between bvFTD and Controls at baseline

| Region | Controls | bvFTD | t-value | p-value |
| --- | --- | --- | --- | --- |
| Lateral Ventricles | 1.87±0.82 | 3.2±0.98 | *11.24* | <0.001 |
| Third Ventricle | 1.4±0.35 | 1.91±0.4 | *10.75* | <0.001 |
| Thalamus-R | 0.9±0.07 | 0.8±0.07 | *-10.61* | <0.001 |
| Amygdala-L | 0.99±0.08 | 0.83±0.13 | *-9.80* | <0.001 |
| Thalamus-L | 0.88±0.07 | 0.79±0.07 | *-9.77* | <0.001 |
| Lateral Orbitofrontal-L | 1±0.07 | 0.89±0.11 | *-9.71* | <0.001 |
| Putamen-R | 0.89±0.08 | 0.76±0.11 | *-9.51* | <0.001 |
| Superior Frontal-R | 0.94±0.08 | 0.83±0.1 | *-9.36* | <0.001 |
| Insula-L | 0.96±0.07 | 0.85±0.11 | *-8.87* | <0.001 |
| Putamen-L | 0.9±0.09 | 0.77±0.11 | *-8.56* | <0.001 |
| Ventral Diencephalon | 0.95±0.06 | 0.87±0.07 | *-8.47* | <0.001 |
| Accumbens-L | 0.9±0.12 | 0.76±0.12 | *-8.37* | <0.001 |
| Amygdala-R | 1±0.09 | 0.87±0.13 | *-8.35* | <0.001 |
| Insula-R | 0.95±0.07 | 0.85±0.11 | *-8.30* | <0.001 |
| Superior Frontal-L | 0.95±0.07 | 0.86±0.11 | *-8.00* | <0.001 |
| Lateral Orbitofrontal-R | 0.99±0.07 | 0.9±0.12 | *-7.93* | <0.001 |
| Accumbens-R | 0.84±0.12 | 0.7±0.09 | *-7.92* | <0.001 |
| Inferior Lateral Ventricle-R | 1.1±0.21 | 1.48±0.52 | *7.63* | <0.001 |
| Pallidum-R | 0.85±0.09 | 0.74±0.11 | *-7.34* | <0.001 |
| Inferior Lateral Ventricle-L | 1.02±0.17 | 1.32±0.41 | *7.33* | <0.001 |
| Pallidum-L | 0.87±0.1 | 0.76±0.13 | *-7.05* | <0.001 |
| Middle Temporal-L | 0.96±0.1 | 0.87±0.1 | *-6.80* | <0.001 |
| Entorhinal-L | 1.13±0.14 | 0.98±0.16 | *-6.66* | <0.001 |
| Middle Temporal-R | 1.01±0.1 | 0.92±0.11 | *-6.47* | <0.001 |
| Medial Orbitofrontal-R | 1±0.09 | 0.9±0.13 | *-6.40* | <0.001 |
| Medial Orbitofrontal-L | 0.93±0.09 | 0.84±0.12 | *-6.27* | <0.001 |
| Entorhinal-R | 1.15±0.18 | 1.01±0.18 | *-6.00* | <0.001 |
| Inferior Temporal-R | 1.08±0.11 | 0.99±0.13 | *-5.61* | <0.001 |
| Caudal Middle Frontal-R | 0.96±0.14 | 0.84±0.15 | *-5.52* | <0.001 |
| Pars Orbitalis-R | 1.05±0.19 | 0.92±0.2 | *-5.49* | <0.001 |
| Rostral Middle Frontal-R | 0.99±0.12 | 0.91±0.15 | *-5.31* | <0.001 |
| Pars Opercularis-L | 0.98±0.15 | 0.86±0.17 | *-4.78* | <0.001 |
| Brain Stem | 1.02±0.08 | 0.96±0.07 | *-4.70* | <0.001 |
| Pars Triangularis-R | 1.03±0.18 | 0.9±0.21 | *-4.68* | <0.001 |
| Caudal Middle Frontal-L | 1.01±0.17 | 0.89±0.17 | *-4.62* | <0.001 |
| Inferior Temporal-L | 1.08±0.12 | 1±0.14 | *-4.51* | <0.001 |
| Inferior Parietal-L | 1.02±0.13 | 0.94±0.13 | *-4.19* | <0.001 |
| Isthmus Cingulate-R | 0.93±0.13 | 0.86±0.12 | *-4.17* | <0.001 |
| Rostral Middle Frontal-L | 0.94±0.11 | 0.88±0.13 | *-4.15* | <0.001 |
| Cerebellum-L | 0.98±0.08 | 0.92±0.07 | *-4.04* | <0.001 |
| Pars Triangularis-L | 1.02±0.17 | 0.94±0.17 | *-4.02* | <0.001 |
| Cerebellum-R | 0.98±0.08 | 0.93±0.08 | *-3.82* | <0.001 |
| Precuneus-R | 0.93±0.09 | 0.89±0.11 | *-3.71* | <0.001 |
| Basal Forebrain | 0.94±0.07 | 0.89±0.07 | *-3.68* | <0.001 |
| Isthmus Cingulate-L | 0.96±0.12 | 0.9±0.14 | *-3.59* | <0.001 |
| Inferior Parietal-R | 0.96±0.12 | 0.9±0.12 | *-3.52* | 0.001 |
| Pars Opercularis-R | 0.98±0.15 | 0.9±0.17 | *-3.51* | 0.001 |
| Hippocampus-R | 0.99±0.09 | 0.93±0.11 | *-3.46* | 0.001 |
| Precuneus-L | 0.92±0.09 | 0.88±0.09 | *-3.31* | 0.001 |
| Para hippocampal-R | 1.01±0.12 | 0.96±0.11 | *-3.28* | 0.002 |
| Supramarginal-R | 1.03±0.14 | 0.96±0.15 | *-3.24* | 0.002 |
| Caudate-R | 0.94±0.09 | 0.87±0.14 | *-3.04* | 0.004 |
| Fusiform-R | 1.01±0.11 | 0.96±0.11 | *-2.77* | 0.009 |
| Hippocampus-L | 1±0.09 | 0.96±0.11 | *-2.77* | 0.009 |
| Superior Parietal-R | 1.01±0.13 | 0.96±0.14 | *-2.62* | 0.014 |
| Superior Temporal-R | 1.04±0.09 | 1.01±0.1 | *-2.58* | 0.016 |
| Pars Orbitalis-L | 1.09±0.18 | 1.02±0.21 | *-2.52* | 0.019 |
| Para hippocampal-L | 1.03±0.12 | 0.99±0.12 | *-2.48* | 0.020 |
| Precentral-L | 1.04±0.1 | 1±0.11 | *-2.45* | 0.022 |
| Vermis | 0.96±0.08 | 0.93±0.09 | *-2.32* | 0.030 |
| Precentral-R | 1.04±0.08 | 1±0.12 | *-2.21* | 0.039 |
| Superior Temporal-L | 0.99±0.09 | 0.95±0.1 | *-2.13* | 0.047 |
| Rostral Anterior Cingulate-L | 0.97±0.19 | 0.93±0.18 | *-1.59* | 0.151 |
| Rostral Anterior Cingulate-R | 0.95±0.18 | 0.91±0.17 | *-1.34* | 0.241 |
| Caudate-L | 0.92±0.09 | 0.89±0.14 | *-1.33* | 0.241 |
| Supramarginal-K | 0.99±0.14 | 0.97±0.17 | *-1.31* | 0.246 |
| Posterior Cingulate-R | 0.96±0.1 | 0.93±0.13 | *-1.23* | 0.277 |
| Lateral Occipital-R | 1.05±0.14 | 1.03±0.1 | *-1.21* | 0.287 |
| Postcentral-L | 1.07±0.11 | 1.05±0.12 | *-1.11* | 0.333 |
| Lingual-L | 1±0.11 | 1.02±0.12 | *1.04* | 0.365 |
| Posterior Cingulate-L | 0.91±0.1 | 0.9±0.13 | *-0.93* | 0.424 |
| Transverse Temporal-L | 1.15±0.26 | 1.15±0.22 | *-0.88* | 0.448 |
| Cuneus-R | 1.06±0.13 | 1.09±0.15 | *0.79* | 0.501 |
| Superior Parietal-L | 1±0.12 | 0.97+0.13 | *-0.74* | 0.527 |
| Lateral Occipital-L | 1.15±0.13 | 1.14±0.13 | *-0.52* | 0.675 |
| Cuneus-L | 0.98±0.13 | 0.97±0.13 | *-0.53* | 0.675 |
| Fourth Ventricle | 1.03±0.32 | 1.04±0.25 | *0.49* | 0.688 |
| Lingual-R | 1.05±0.1 | 1.06±0.11 | *0.47* | 0.694 |
| Paracentral-R | 0.93±0.13 | 0.91±0.16 | *-0.47* | 0.694 |
| Fusiform-L | 0.97±0.11 | 0.96±0.12 | *-0.42* | 0.722 |
| Paracentral-L | 0.95±0.13 | 0.93±0.14 | *0.34* | 0.771 |
| Caudal Anterior Cingulate-L | 0.9±0.19 | 0.9±0.16 | *-0.29* | 0.805 |
| Caudal Anterior Cingulate-R | 1.03±0.2 | 1.02±0.19 | *0.28* | 0.806 |
| Pericalcarine-L | 0.94±0.15 | 0.94±0.16 | *-0.23* | 0.840 |
| Pericalcarine-R | 0.97±0.15 | 0.96±0.16 | *0.15* | 0.893 |
| Transverse Temporal-R | 0.87±0.16 | 0.87±0.16 | *-0.04* | 0.978 |
| Postcentral-R | 1.02±0.1 | 1.03±0.13 | *0.00* | 0.998 |

Model: DBM ~ 1 + Dx + AGE + Dx:AGE + SEX + (1|ID) + (1|SITE). Values expressed as mean DBM Jacobian value ± SD. Note that the *mean DBM Jacobian* is a multiplicative factor: for example, lateral ventricles in the Controls are 1.87 times larger than the lateral ventricles of the ICBM152 template volume created from young adults. Negative t-values indicate atrophy in patients with bvFTD compared with Controls; positive t-values indicate enlargement in bvFTD relative to controls. L= left, R=right.

**Table S3:** Annual change in DBM per region and sample size estimation both cohorts

| Region | Controls  (ΔDBM Jacobian) | bvFTD  (ΔDBM Jacobian) | p-Value | Sample Size  12 months | Sample Size  24 months |
| --- | --- | --- | --- | --- | --- |
| Lateral Ventricles | 0.06±0.06 | 0.34±0.32 | <0.001 | 194 | 50 |
| Posterior Cingulate-L | 0±0.01 | -0.02±0.03 | <0.001 | 251 | 65 |
| Ventral Diencephalon | 0±0.01 | -0.02±0.02 | <0.001 | 257 | 66 |
| Third Ventricle | 0.03±0.03 | 0.12±0.13 | <0.001 | 264 | 68 |
| Putamen-L | 0±0.01 | -0.03±0.03 | <0.001 | 282 | 73 |
| Putamen-R | 0±0.01 | -0.03±0.03 | <0.001 | 286 | 73 |
| Isthmus Cingulate-L | 0±0.03 | -0.02±0.03 | 0.001 | 306 | 79 |
| Inferior Lateral Ventricle-L | 0.01 | 0.1±0.14 | <0.001 | 344 | 89 |
| Middle Temporal-R | 0±0.02 | -0.02±0.03 | <0.001 | 433 | 110 |
| Thalamus-L | 0±0.01 | -0.02±0.02 | <0.001 | 460 | 118 |
| Middle Temporal-L | 0±0.02 | -0.02±0.03 | 0.001 | 468 | 119 |
| Superior Temporal-L | 0±0.01 | -0.02±0.03 | <0.001 | 475 | 121 |
| Superior Parietal-R | 0.01±0.06 | -0.02±0.04 | 0.019 | 493 | 126 |
| Inferior Lateral Ventricle-R | 0.01 | 0.1±0.18 | <0.001 | 542 | 138 |
| Precuneus-L | 0±0.02 | -0.01±0.03 | 0.001 | 570 | 145 |
| Caudate-L | 0‑0.02 | 0.04±0.;09 | <0.001 | 586 | 149 |
| Amygdala-L | -0.01±0.02 | -0.04±0.07 | <0.001 | 593 | 150 |
| Thalamus-R | 0±0.01 | -0.01±0.02 | <0.001 | 605 | 154 |
| Pallidum-R | 0±0.02 | -0.02±0.05 | <0.001 | 605 | 154 |
| Basal Forebrain | 0±0.02 | -0.02±0.04 | 0.001 | 620 | 157 |
| Brain Stem | 0±0.01 | -0.01±0.02 | <0.001 | 635 | 161 |
| Inferior Temporal-L | 0±0.02 | -0.02±0.04 | 0.005 | 715 | 181 |
| Posterior Cingulate-R | 0±0.02 | -0.02±0.04 | 0.001 | 725 | 184 |
| Insula-L | 0±0.01 | -0.01±0.02 | 0.001 | 742 | 187 |
| Supramarginal-K | 0±0.03 | -0.03±0.07 | 0.001 | 745 | 188 |
| Caudal Middle Frontal-L | 0±0.03 | -0.02±0.05 | 0.003 | 755 | 191 |
| Fusiform-L | 0±0.02 | -0.01±0.03 | 0.003 | 821 | 208 |
| Pallidum-L | 0±0.03 | -0.02±0.05 | 0.005 | 822 | 208 |
| Para hippocampal-L | 0±0.02 | -0.01±0.04 | 0.003 | 840 | 212 |
| Para hippocampal-R | 0±0.02 | -0.02±0.04 | 0.002 | 918 | 232 |
| Superior Temporal-R | 0±0.02 | -0.01±0.03 | 0.013 | 968 | 245 |
| Inferior Temporal-R | 0±0.02 | -0.02±0.05 | 0.005 | 1045 | 263 |
| Pars Orbitalis-L | 0±0.05 | -0.03±0.1 | 0.008 | 1074 | 270 |
| Caudate-R | 0±0.02 | 0.02±0.07 | 0.002 | 1172 | 295 |
| Rostral Middle Frontal-L | 0±0.03 | -0.02±0.06 | 0.015 | 1187 | 299 |
| Pars Opercularis-L | 0±0.02 | -0.02±0.05 | 0.007 | 1220 | 307 |
| Superior Frontal-R | 0±0.02 | -0.01±0.03 | 0.033 | 1229 | 310 |
| Precentral-L | 0±0.02 | -0.01±0.03 | 0.033 | 1315 | 331 |
| Cerebellum-R | 0±0.01 | -0.01±0.02 | 0.008 | 1398 | 352 |
| Precuneus-R | 0±0.03 | -0.01±0.04 | 0.049 | 1459 | 367 |
| Amygdala-R | -0.01±0.03 | -0.03±0.06 | 0.017 | 1500 | 377 |
| Superior Frontal-L | 0±0.01 | -0.01±0.03 | 0.014 | 1612 | 406 |
| Entorhinal-L | 0±0.03 | -0.03±0.09 | 0.017 | 1650 | 414 |
| Medial Orbitofrontal-R | 0±0.02 | -0.02±0.04 | 0.035 | 1663 | 418 |
| Paracentral-R | 0±0.04 | -0.01±0.06 | 0.037 | 1676 | 421 |
| Fourth Ventricle | 0±0.02 | 0.02±0.06 | 0.022 | 1777 | 446 |
| Medial Orbitofrontal-L | -0.01±0.02 | -0.02±0.04 | 0.047 | 1871 | 470 |
| Supramarginal-R | 0±0.04 | -0.03±0.11 | 0.005 | 1896 | 476 |

Table expresses mean yearly change in DBM Jacobian value ± SD, p-Value regressed by sex and age and FDR corrected for multiple comparisons and sample size needed in order to reach 25% reduction in the rate of atrophy/enlargement per region in bvFTD patients. All estimates were multiplied by 1.2 to account for the expected attrition. L= left, R=right.

**Table S4:** Annual change in clinical scores for bvFTD vs Controls

|  | Controls | bvFTD | p Value | Sample Size |
| --- | --- | --- | --- | --- |
| CDR-SOB | -0.03±0,31 | 2.54±2.97 | <0.001 | 204 |
| MMSE | 0.06±1.11 | -2.76±5.39 | <0.001 | 552 |
| MTT time | -6.47±17.96 | -6.72±29.4 | >0.9 | 2142746 |
| Semantic Fluency | -0.41±4.38 | -1.94±5.51 | 0.078 | 1946 |
| Letter Fluency | 0.23±4.52 | -1.12±3.38 | 0.154 | 945 |
| MOCA | 0.66±2.71 | -2.18±4.06 | 0.003 | 310 |
| CGI | -0.03±0.15 | 0.36±0.86 | 0.008 | 720 |

Table expresses mean yearly change in functional/cognitive scores ± SD, p-Value regressed by sex and age and FDR corrected for multiple comparisons and sample size needed in order to reach 25% reduction in the rate of atrophy/enlargement per region in bvFTD patients. All estimates were multiplied by 1.2 to account for the expected attrition.
